# Immediate cortical responses to transcranial magnetic stimulation of the human motor leg area

**DOI:** 10.64898/2026.09.30.755678

**Authors:** Mikkel Malling Beck, Marten Nuyts, Laura Sakalauskaite, John Rothwell, Hartwig Roman Siebner, Lasse Christiansen

## Abstract

**Background:** Immediate transcranial evoked potentials (iTEPs) are ultra-early electroencephalographic responses elicited by single-pulse transcranial magnetic stimulation (TMS) of the primary motor cortical hand area (M1-HAND). Their characteristic latency, waveform, and sensitivity to short-interval intracortical inhibition suggest a close relationship to the cortical generators of indirect corticospinal volleys (I-waves). Here, we tested the hypothesis that iTEPs constitute a fundamental physiological response of human motor cortex and therefore generalize across motor representations, including the leg representation (M1-LEG).

**Methods:** Single-pulse TMS was delivered over the motor cortical representation of the right tibialis anterior (TA) in 16 healthy volunteers. In a subset of participants (n=10), stimulus–response curves and paired-pulse short-interval intracortical inhibition (SICI) measures were obtained.

**Results:** TMS of M1-LEG elicited reproducible iTEPs consisting of a slow positive deflection with superimposed high-frequency peaks with topographic maxima in electrodes adjacent to the stimulation site. M1-LEG iTEPs emerged close to active motor threshold and increased in prevalence and amplitude with increasing stimulation intensity. Paired-pulse stimulation produced consistent SICI of the second and third iTEP peaks. M1-LEG stimulation also evoked conventional later TEPs with latencies and polarities closely resembling those recorded after M1-HAND stimulation.

**Conclusions:** TMS of the human M1-LEG elicits robust iTEPs that closely resemble those recorded from M1-HAND in their timing, intensity dependence, and sensitivity to intracortical inhibition. These findings indicate that iTEPs are not specific to M1-HAND but reflect a general physiological response of human motor cortex closely linked to corticospinal I-wave generation. This broadens their utility as a non-invasive probe of excitatory and inhibitory circuit dynamics throughout the human motor system.

**Highlights:**

- Single-pulse TMS of the leg motor cortex (M1-LEG) elicits reproducible immediate TMS-evoked potentials (iTEPs).
- M1-LEG iTEPs emerge near active motor threshold, increase with stimulation intensity, and display selective short-interval intracortical inhibition of late iTEP peaks with paired-pulse TMS.
- Immediate and later TMS-evoked responses show striking resemblance in leg and hand motor cortex.
- The shared properties of M1-LEG and M1-HAND iTEPs support iTEPs as a general motor cortex response linked to corticospinal I-wave generation and intracortical inhibition.

## Introduction

Transcranial magnetic stimulation (TMS) enables non-invasive probing of human brain function in real time by inducing transient neural activity and assessing the resulting physiological responses (Siebner et al., 2022). The effects of TMS on the primary motor cortex (M1) are most commonly quantified using motor evoked potentials (MEPs), which can be readily recorded from skeletal muscles using surface electromyography (EMG). Combining TMS with electroencephalography (EEG) enables direct assessment of local cortical reactivity and network responses through TMS-evoked EEG potentials (TEPs)(Beck et al., 2024b; Hernandez-Pavon et al., 2023). Single-pulse stimulation of the hand representation of the primary motor cortex (M1-HAND) elicits a characteristic sequence of TEP components, including the recently described immediate TEP (iTEP)(Beck et al., 2024a), which is spatially confined to the pericentral cortex (Nuyts et al., 2025). Subsequent early TEP components (∼10-70 ms) are thought to reflect reverberating activity within local cortical circuits as well as through cortico-subcortical and cortico-cortical reentrant loops, whereas later components (∼70 ms and later) show less spatial and temporal specificity across stimulation sites (Conde et al., 2019). To date, TMS-EEG studies of the motor cortex have predominantly focused on lateral M1 regions controlling upper extremity muscles, particularly the intrinsic hand muscles (Beck et al., 2024b). In contrast, it remains unknown whether stimulation of the medial M1 containing the leg motor representation (M1-LEG) elicits comparable cortical responses.

The human M1 is unique compared to other cortical regions exhibiting several distinctive anatomical and microstructural features, including giant Betz cells, dense cortical myelination, a thin or absent granular layer IV, and higher supra- vs infragranular receptors (Palomero-Gallagher and Zilles, 2019). The M1-LEG is located predominantly within the anterior paracentral lobule along the banks of the interhemispheric fissure (Sato et al., 2022; Terao et al., 1998). This region contains Betz cells (Rivara et al., 2003) and displays cytoarchitecture and microstructural characteristics comparable to those of M1-HAND (Fukutomi et al., 2018). Consistent with these structural similarities, M1-HAND and M1-LEG have other physiological similarities. They share neurochemical and oscillatory properties (Hansen et al., 2022), exhibit parallel somatotopically organized functional connectivity with subcortical motor structures (Zeharia et al., 2015), and show highly comparable patterns of intracortical inhibition and facilitation when assessed using conventional MEP-based paired-pulse TMS paradigms (Mrachacz-Kersting et al., 2021; Stokić et al., 1997).

Beyond these shared cortical properties, electrophysiological studies further demonstrate that, like upper-extremity muscles, leg muscles are controlled through mono-, di-, and polysynaptic corticospinal pathways linking pyramidal tract projection neurons to spinal motoneurons (Cowan et al., 1986; Maertens De Noordhout et al., 1999; Nielsen and Petersen, 1995). Notably, fast monosynaptic corticospinal projections are thought to contribute to the rapid and precise execution of leg movements (Nielsen and Petersen, 1995). Taken together, the substantial structural, neurophysiological, and connectivity-related similarities between M1-LEG and M1-HAND suggest that TMS of the M1-LEG is expected to evoke immediate, as well as early and late-latency TEP responses, analogous to those observed following M1-HAND stimulation.

Invasive epidural recordings obtained at cervical as well as upper and lower thoracic levels have demonstrated that single-pulse TMS over M1-HAND and M1-LEG elicits a descending corticospinal volley consisting of multiple waves (Burke et al., 1993; Di Lazzaro et al., 1998a; V. Di Lazzaro et al., 2001). These comprise both a direct wave (D-wave), suggested to reflect axonal activation, and multiple indirect waves (I-waves), thought to arise from transsynaptic activation of large-diameter pyramidal tract neurons. Since individual corticospinal neurons typically discharge only a single action potential in response to a TMS pulse, the characteristic periodicity of the I-waves (approximately 500–1000 Hz) is thought to reflect the synchronized recruitment of neuronal populations rather than repetitive firing of individual neurons (Yu et al., 2024). The presence of descending corticospinal volleys in thoracic epidural recordings following M1-LEG stimulation, together with the periodic peaks observed in post-stimulus time histograms recorded from the tibialis anterior (TA) muscle (Brouwer and Ashby, 1990), suggests that lower-extremity muscles may be subject to direct cortical control through mechanisms comparable to those described for upper-extremity muscles. However, direct characterization of these corticospinal volleys remains limited because it relies on invasive epidural recordings.

Recent advances in TMS-EEG have enabled characterization of the immediate cortical effects of TMS within the first 2-10 ms after single-pulse stimulation, enabling characterization of both the immediate local cortical response pattern as well as the subsequent propagation and reverberatory activity within cortico-subcortical and cortico-cortical networks (Beck et al., 2024a). Leveraging these methodological advances in TMS-EEG, the present study aimed to determine whether TEP responses over M1-LEG (1) exhibit physiological characteristics analogous to those observed for M1-HAND, (2) are dose-dependent, and (3) are sensitive to paired-pulse inhibition. Our overarching objective was to establish a non-invasive electrophysiological framework for investigating immediate cortical responses associated with corticofugal signaling and distributed brain network activity evoked by focal TMS targeting the leg representation of the human M1.

## Methods

### Participants

Twenty healthy volunteers without neurological and psychiatric disorders or contraindications to TMS or MRI participated in the experiment after giving informed consent. Participants were excluded if it was not possible to record EEG responses to TMS of the M1-LEG free of clear scalp muscle artefacts at 110% of resting motor threshold (RMT) after a personalized TEP optimization procedure (see below for further details). Three subjects were excluded on this basis. A fourth subject was excluded due to excessive eye blinks (more than 50% of trials) leaving 16 participants from whom data is reported in this manuscript. The experiments were carried out in accordance with the Helsinki Declaration and were approved by the regional ethical committee (Capital Region of Denmark; Protocol number H-15008824).

### Experimental procedure

To characterize the cortical response to TMS of the M1-LEG, participants took part in a single experimental session in which single- and paired-pulse TMS were delivered at different intensities to the motor cortical leg representation while their EEG was recorded. The right TA was targeted across both the main and sub-experiments. For the main experiment, stimulation was delivered to the M1-LEG at intensities corresponding to 95% and 110% of the right TA’s resting motor threshold (RMT) (main experiment; Figure 1A). These intensities were chosen, as we have previously shown that prominent EEG responses are elicited from the M1-HAND with these same intensities. Additionally, a subset of participants (10/16) underwent a detailed stimulus-response mapping and a paired-pulse conditioning protocol to investigate how evoked cortical responses were contingent on stimulation intensity, and how they were influenced by a short-interval intracortical inhibition (SICI) paired pulse paradigm (sub-experiments, Figure 1B). The main experiment was always carried out first followed by the stimulus-response experiment and finally the SICI experiment. The order of conditions within each of the experiments was randomized. Across all conditions, data was recorded in blocks of 100 stimulations.

**Figure 1.**
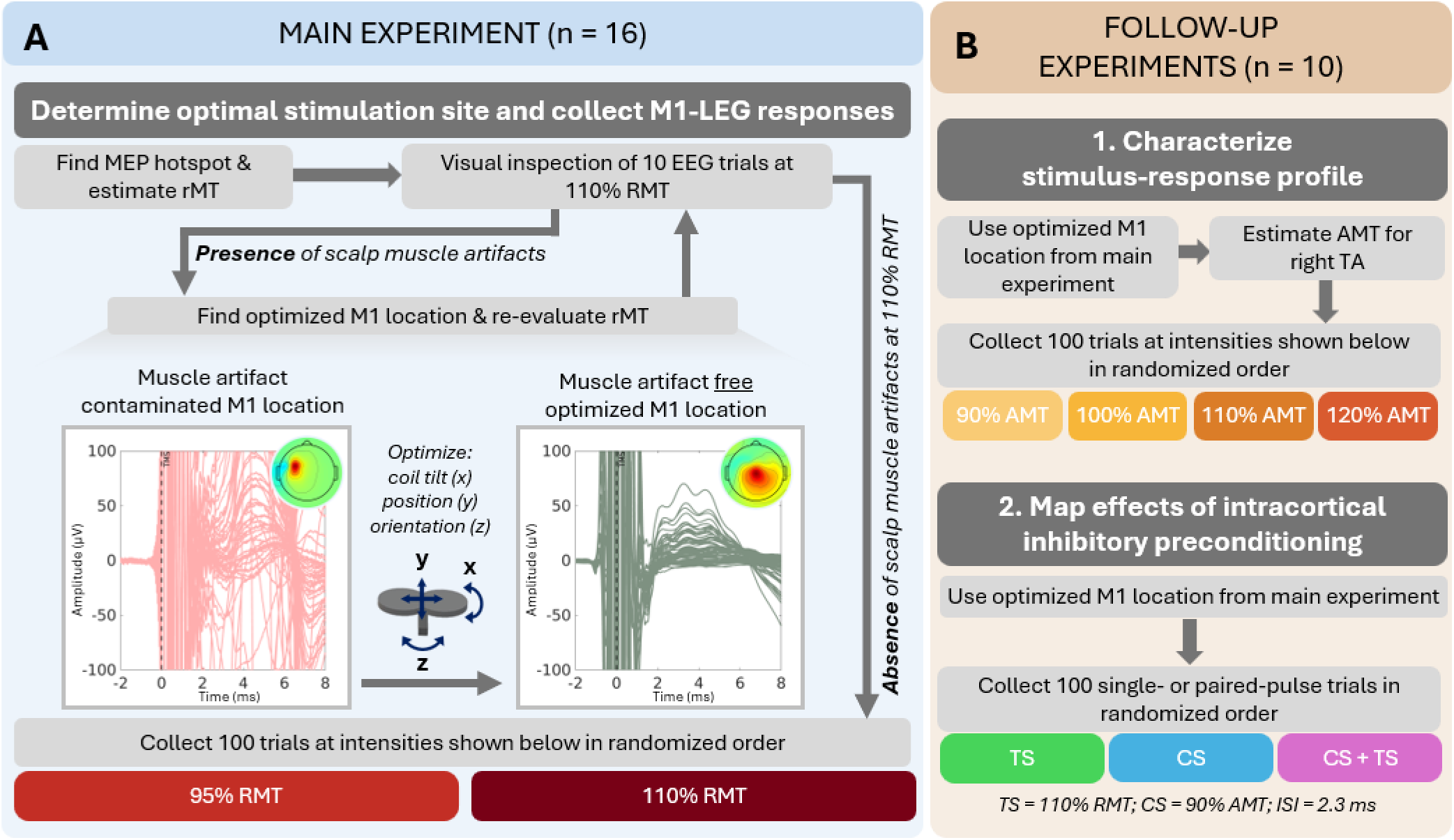
Experimental procedure. Participants (n=16) underwent a main experiment (A) and a subset (n=10) also participated in two sub-experiments (B). The experimental procedure for each of the experimental conditions is outlined. As part of the main experiment, the optimal position for stimulating the M1-LEG without evoking scalp muscle artefacts was determined. In short, the coil position was optimized to elicit MEPs in *m. tibialis anterior* concomitantly with artifact-free immediate TEPs in the EEG. If muscle artefacts were present, the coil was tilted or rotated to direct the induced electric field away from the *m. temporalis or m. frontalis*. Then, M1-LEG was stimulated at 95% and 110% RMT. In the sub-experiments, a detailed mapping of the stimulus-response profile was determined by stimulating at intensities from 90% to 120% AMT at the optimal site identified in the main experiment. Subsequently, the Kujirai short-interval intracortical inhibition (SICI) paired pulse TMS paradigm (Kujirai et al., 1993) was adopted to study intracortical inhibition of iTEPs.

For all experimental conditions, participants were sitting at rest with their eyes fixated at a cross in front of them and instructed to avoid blinking excessively. Participants were equipped with modified earplugs playing an audio file generated using TAAC software consisting of pink noise overlaid with recorded TMS clicks (Russo et al., 2022). The sound pressure was adjusted until participants reported that they were unable to hear the coil discharging or at the highest tolerable level that attenuated the TMS “click” sound considerably. A thin layer of foam (approx. 1.5 mm thick) was placed between the coil and the electrodes to reduce bone conduction of sound and vibration-induced somatosensory co-activation.

### Transcranial magnetic stimulation

Biphasic TMS pulses (direction: AP-PA; duration: 280µS) were delivered every 2-sec (± 10% jitter) through a 70-mm figure-of-eight coil (MC-B70; MagVenture A/S, Farum, Denmark) using a MagVenture X100 stimulator (MagVenture A/S, Farum, Denmark). The recharge delay was set at 500 ms after stimulation to avoid recharge artifacts in the EEG recordings. Stimulations were controlled using a data acquisition unit (Micro1401-4, Cambridge Electronic Design Ltd., Cambridge, UK) using Signal software (v4.17, Cambridge Electronic Design Ltd., Cambridge, UK).

Stimulations were targeting the M1-LEG area. Specifically, the M1-LEG stimulation site was determined based on two criteria in each participant: (1) it should effectively cause MEPs in the right TA muscle (2) it should not cause scalp muscle activation.

To identify the initial stimulation spot, the coil was initially placed at the vertex (Cz) with the coil handle pointing backwards. A mini-mapping procedure was then performed to localize the right TA motor hotspot by moving or rotating the coil while the presence of MEPs was visually confirmed. Motor hotspots for lower extremity muscles are known to be more variable across subjects compared to that of e.g. the intrinsic hand muscles. In 4/16 volunteers, the hotspot was found by rotating the coil (90 deg. to the midline) and reverting the current direction electronically in the magnetic stimulator, such that the dominant (second) phase of the biphasic pulse induced a current in the left hemisphere. The TMS coil position was monitored via stereotaxic neuronavigation (TMS navigator, Localite, GmhB, Bonn, Germany) using individual T1w images (3T MRI, Siemens Prisma or Philips Achieva).

After having found the TA motor hotspot, EEG responses to TMS were inspected for the presence of scalp muscle artifacts using a procedure that we have previously described for stimulation of the M1-HAND (Beck et al., 2024a)(Figure 1A). In short, 10-20 TMS pulses were delivered at the highest target intensity (110% resting motor threshold; RMT) and responses in the EEG leads were visually inspected for the presence of scalp muscle artefacts. The following criteria were adopted to determine the presence of scalp muscle artefacts: (1) high-amplitude responses (>100µV); (2) bipolar waveforms resembling compound muscle action potentials; (3) topographical maxima in temporal and/or frontal electrodes overlying *m. temporalis* and *m. frontalis* (Figure 1A). If scalp muscle artefacts were present (9/16 individuals at motor hotspot) minor corrections of the coil position, tilt, and/or rotation were performed away from the scalp muscles activated (*m. temporalis* and/or *m. frontalis*) until absence of scalp muscle activation was confirmed (Figure 1A). The new stimulation site was saved, a new RMT was estimated, and the procedure outlined above was repeated until no scalp muscle co-activation was observed.

In the main experiment, stimulation was delivered at 95% and 110% RMT over the optimized M1-LEG stimulation site. The RMT was estimated as the lowest intensity leading to MEPs in the right TA muscle of 50µV in 5/10 consecutive trials. The average RMT and standard deviation across participants were 66.8% ± 10 of maximal stimulator output (MSO) at the optimized stimulation site.

For the stimulus-response protocol, participants received TMS pulses at 90%, 100%, 110% and 120% of active motor threshold (AMT). The AMT was determined as the lowest intensity that yielded clear MEPs in 5/10 consecutive trials during a sustained isometric contraction of the right TA muscle at ∼ 10% maximal voluntary contraction. The average AMT and standard deviation across participants were 48.6% ± 4.3 MSO. For the SICI protocol, an interstimulus interval of 2.3ms was used and the intensity of the conditioning pulse and the test pulse was set at 90% AMT and 110% RMT, respectively (Kujirai et al., 1993).

### EEG and EMG recordings

EEG was recorded with BrainVision Recorder software (Brain Prdoucts, GmhB, Germany) from 61 passive Ag/AgCl electrodes placed in an equidistant EEG cap (M10 cap layout, BrainCap TMS, Brain Products, GmbH, Germany). The reference and the ground were placed on the right and left side of the forehead, respectively. Electrodes were prepared using electroconductive and abrasive gel to reduce impedance levels (≤5kΩ). Impedance levels were checked throughout the experiment. A TMS-compatible EEG amplifier (ActiCHamp Plus 64 System, Brain Products, Germany) was used at a sampling rate of 50 kHz (anti-aliasing at 10.3 kHz) as this has shown to be capable of capturing EEG responses ∼ 2 ms after stimulation (Beck et al., 2024a).

EMG signals were amplified (x500), bandpass filtered (10-2000Hz), and sampled at 2000 Hz (Digitimer D360, Digitimer Ltd., Hertfordshire, UK) using Signal software (v4.17, Cambridge Electronic Design, Cambridge, UK). EMG responses to TMS were recorded from the right *m. tibialis anterior (TA).* For EMG recordings, the ground was placed on the right patella.

### EEG preprocessing

EEG preprocessing was carried out in MATLAB (v. 2020a) using functions from EEGLAB. In accordance with our previous publications (Beck et al., 2024a; Nuyts et al., 2025), EEG data was epoched from -500 to 500 ms around stimulation and band-pass filtered at 0.1-2000Hz. EEG data was visually inspected for noisy channels and bad epochs. Channels contaminated by line-noise or by decay artefacts after stimulation were removed (median across all conditions: 3 channels). Epochs contaminated by eye-blinks, single-trial residual scalp muscle artefacts, or displaying excessive noise were removed (median across all conditions: 19). Finally, EEG data was baseline-corrected from -110ms to -10ms before stimulation.

### Data analysis and statistics

Following EEG data preprocessing, the global mean field amplitude (GMFA) was computed as a measure of global TMS-evoked cortical activation. Statistical analyses were performed in MATLAB. Akin to our previous publications, we investigated the effects of stimulus intensity over M1-LEG (110% RMT vs. 95% RMT) and paired pulse inhibition (TS vs. CS+TS) using a temporal clustering approach (Christiansen et al., 2026; Nuyts et al., 2025). In brief, a mixed effect model was fitted for each timepoint of the GMFA (dependent variable) in the 2-8ms post-stimulation window comprising the iTEP responses. The mixed models included the stimulus intensity (two levels: 95% RMT vs. 110%RMT) and stimulation type (two levels: TS vs. CS+TS) as the fixed factor with the participant added as the random intercept:

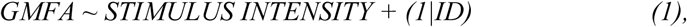

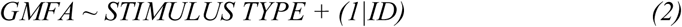

F-values obtained from each of the models outlined were used to calculate the threshold-free clustering enhanced (TFCE) values using the formula proposed in (Smith and Nichols, 2009):

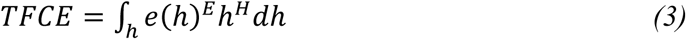

A cluster-based approach was also adopted to investigate the intensity by which the GMFA exceeded baseline activity in the stimulus-response sub-experiment. Specifically, the GMFA time series for each intensity (90% AMT – 120% AMT) was tested against the baseline activity. At each timepoint in the window from 2 to 10 ms, the GMFA was compared to the subject-specific average baseline activity (-10 to -2 ms) using paired t-tests. Statistically significant samples (p < 0.01, two-tailed) were subsequently grouped into temporal clusters, and cluster statistics were computed against a null distribution obtained from 1000 permutations of baseline and post-stimulus data to yield cluster-level p-values.

## Results

### TMS over M1-LEG evokes immediate and classical M1 TEP responses

TMS was delivered over the motor cortical leg representation at 110% RMT in 16 healthy individuals. The evoked EEG responses can be seen in Figure 2. As illustrated in the top panel, a prominent immediate response was observed in the time window from 2-8 ms after stimulation. This response consisted of a series of high-frequency peaks superimposed to a slower positive deflection with topographical maximum close to the site of stimulation. The immediate response was evoked in all individuals but with considerable variation in the magnitude and exact periodicity of individuals response profiles (Figure 2, right panel). This was not directly related to the absolute stimulation intensity (%MSO). Individual peak latencies and amplitudes of the GMFA are reported in supplementary table S1. Latencies and amplitudes were extracted from the GMFA to avoid electrode-specific bias, yielding peak estimates that reliably reflect the global immediate cortical response. In the bottom panel, the evoked responses at a longer timescale can be observed for reference (-50 to 200 ms visualized). This shows a clear pattern consisting of negative deflections around 15 ms, 45 ms and 100 ms alongside positive deflections around 30 ms and 60 ms, resembling the classical M1 TEP response that is also observed after stimulating M1-HAND.

**Figure 2.**
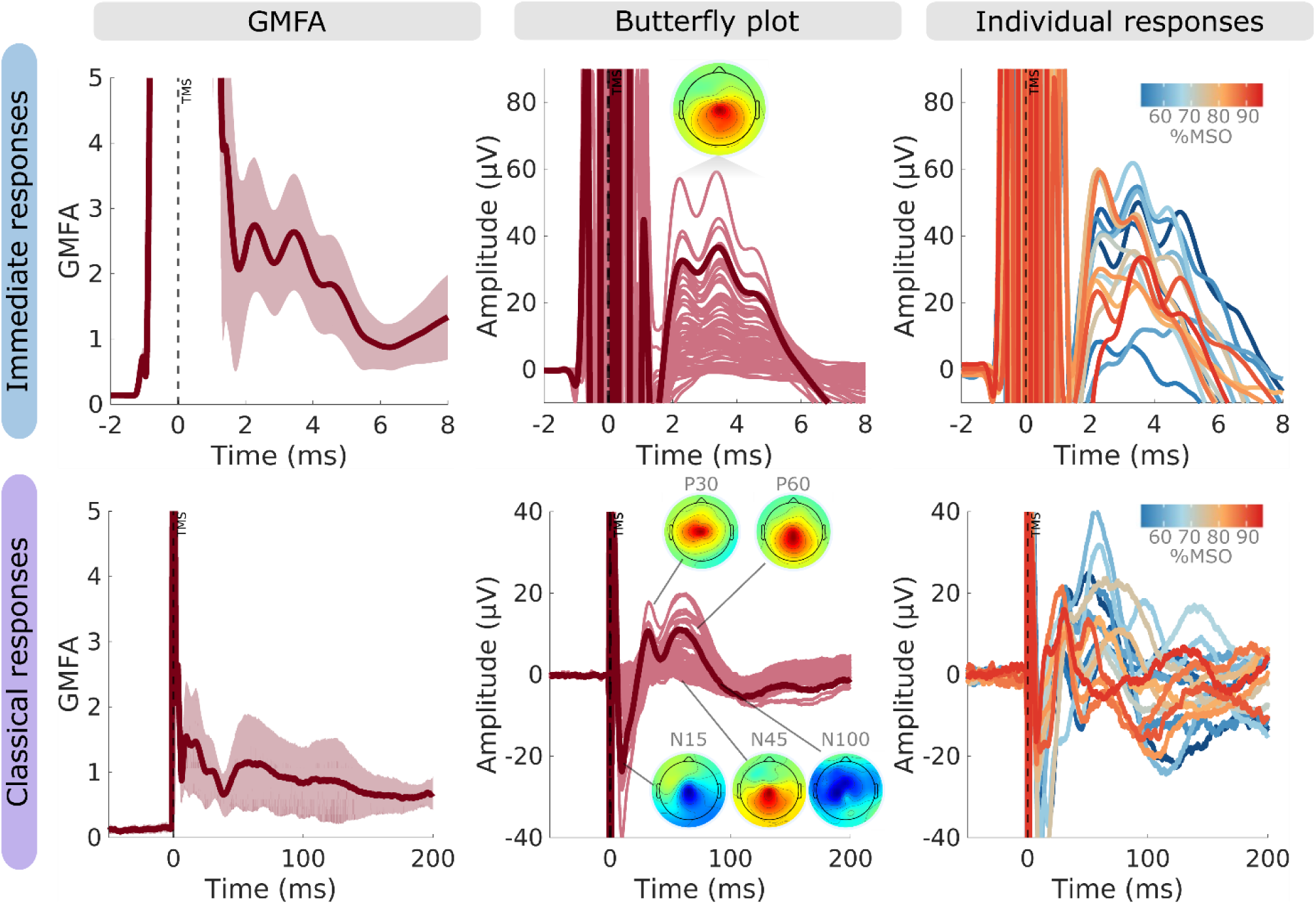
EEG responses evoked from TMS of M1-LEG. The top and bottom row shows the immediate transcranial evoked potential (iTEP) and classical transcranial evoked potential (TEP) elicited from stimulating the M1-LEG at 110% RMT, respectively. Note the different timescales. Group average GMFA +- SD (left; n=16) and butterfly plots with topographic distribution (middle) alongside individual responses (right) to stimulation are depicted. TMS of M1-LEG provokes both immediate and classical TEP responses with a topographical maximum close to the stimulated region. Responses are evoked in all subjects with considerable variability in the latencies and amplitudes. Amplitudes of both immediate and classical responses do not seem to relate to absolute stimulation intensity.

### iTEP responses depend on stimulation intensity

To investigate the effects of stimulation intensity on iTEPs, TMS was delivered at 110% RMT and 95% RMT in 16 individuals. Figure 3A shows that stimulation at 110% RMT produced greater iTEP responses compared to stimulation delivered at 95% RMT. A comparison of the evoked GMFAs revealed a significant difference based on stimulation intensity in the time window from 2.00-8.00ms (Figure 3B). The mean area under the curve (AUC) of the GMFA within the 2.00-8.00 ms window was 76.5 µV*ms after stimulation at 110% RMT and 45.9 µV*ms after stimulation at 95% RMT.

**Figure 3.**
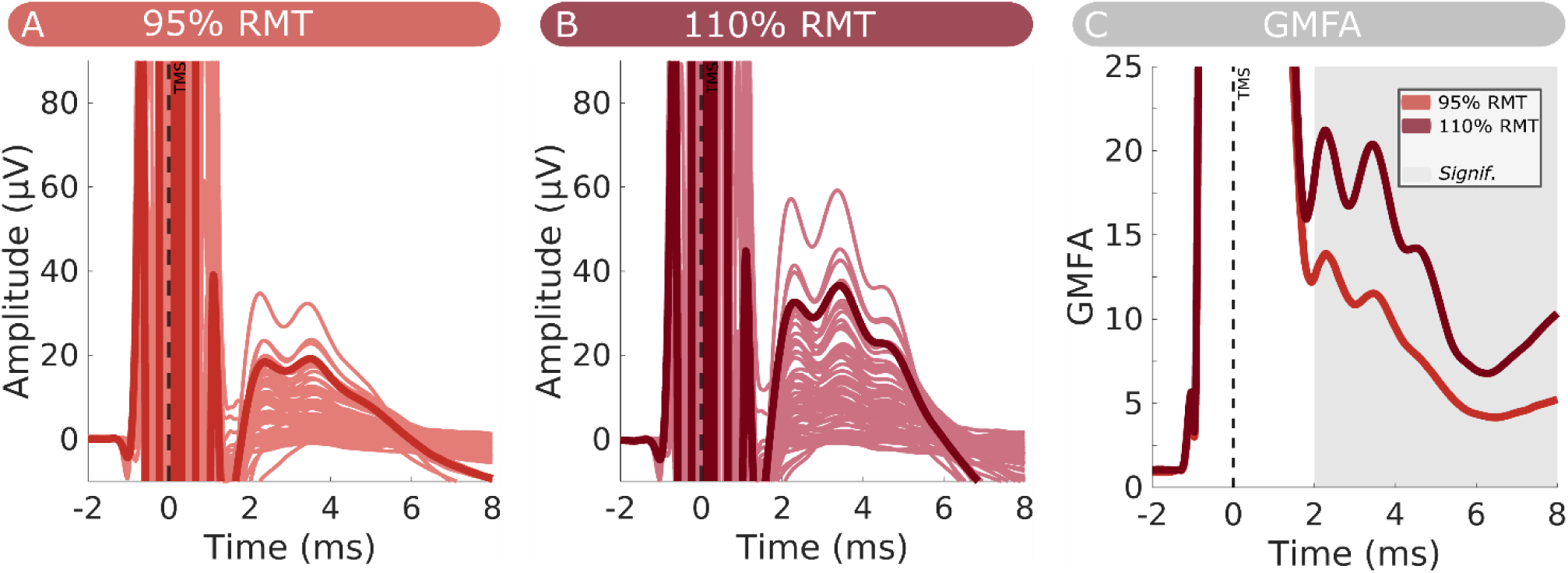
Magnitude of M1-LEG iTEP depend on stimulation intensity. (A) Shows the butterfly plots representing the immediate transcranial evoked potential (iTEP) elicited from stimulating the M1-LEG at 95% and (B) 110% RMT, respectively. The electrode highlighted in the butterfly plot represents an electrode close to the site of stimulation. iTEPs are evoked at both the 95% RMT and 110% RMT intensities, but the response appears greater in magnitude for 110% RMT compared to 95% RMT. (C) Group mean GMFA for the two stimulation intensities. Shaded area represents the time-window of significant differences between stimulating at 95% RMT and 110% RMT. Group data (*n*=16) shown as mean.

To further characterize the iTEP recruitment profile, immediate responses to TMS with intensities extending from 90% of AMT to 120% AMT were recorded in 10 individuals. We compared pre-stimulus baseline activity (-2 to -10 ms) with post-stimulus activity (2 to 10 ms) to investigate the intensity at which reliable iTEPs appeared. This analysis revealed that iTEPs that differed from pre-stimulus baseline activity were observed around 100% AMT (Figure 4). Specifically, when tested up against baseline activity, no significant time-windows were observed between pre-stimulus and post-stimulus GMFA in the 90% AMT condition. In contrast, for 100% AMT, 110% AMT, and 120% AMT significant clusters were observed from 2.00-6.50ms, 2.00-7.70ms, and 2.00-6.87ms, respectively. The mean GMFA for the 2.00-8.00 ms time window was 13.4 µV*ms, 17.7 µV*ms, 20.4 µV*ms and 25.8 µV*ms for the 90% AMT, 100% AMT, 110% AMT, and 120% AMT, respectively. Visually, iTEP response magnitudes also seemed to increase with stimulation intensity and both the first (P1) and second (P2) of the high-frequency iTEP peaks became more prevalent with increasing intensity (Figure 4). When stimulating at 110% AMT and above, a reliable P1 could be extracted from all individuals (10/10). Collectively, this shows that iTEP responses appear around active motor threshold and scale with stimulation intensity.

**Figure 4.**
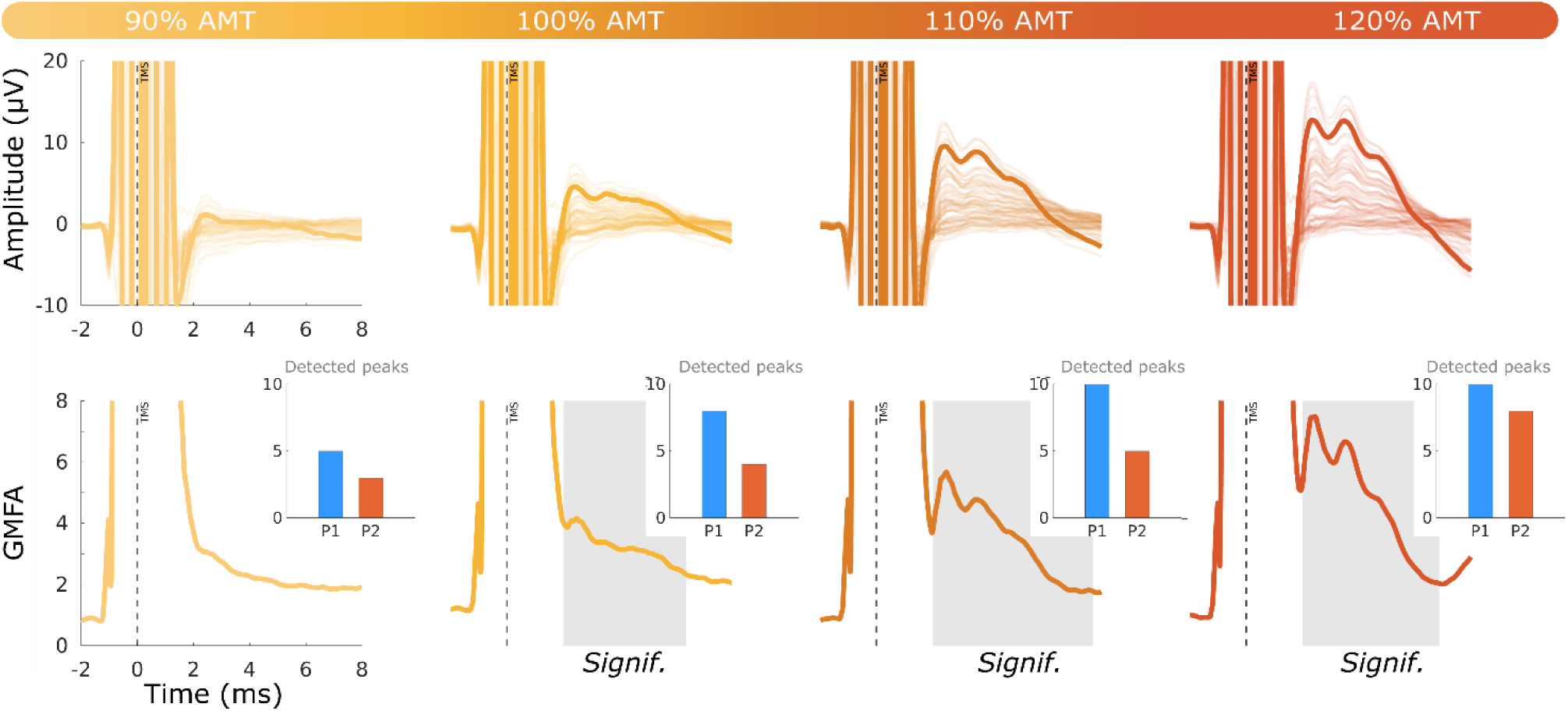
Recruitment profile of M1-LEG iTEPs. Butterfly plots (top) and GMFA (bottom) of immediate responses following stimulation of M1-LEG at intensities ranging from 90% AMT to 120% AMT. The electrode highlighted in the butterfly plot represents an electrode close to the site of stimulation. For the bottom plot of the GMFA, the shaded regions reflect significant clusters where the GMFA between the iTEP window and pre-stimulus baseline activity differs significantly. The insets reflect the proportion of individuals in whom the first (P1) and second (P2) of the high-frequency peaks could be reliably detected in the GMFA. The scaling of the y-axes has been adjusted to account for the overall smaller responses observed at lower intensities. iTEP responses appeared around 100% AMT and increased with stimulation intensity. Group data (*n*=10) shown as mean.

### Short-latency inhibition of immediate TEPs

Ten participants underwent paired pulse TMS following the Kujirai paradigm (Kujirai et al., 1993) to investigate if SICI can observed in iTEPs evoked from M1-LEG. Average MEP inhibition across the 10 individuals verified the expected corticomotor inhibition (Figure 5A). The iTEPs also appeared smaller in magnitude following the CS+TS than for the TS condition (Figure 5B). Cluster-based permutation between the TS and the CS+TS condition revealed a significant difference in the time-window aligning to the 2^nd^ and 3^rd^ iTEP peaks ranging from 2.72ms - 5.08ms as well as from 7.96ms – 8.00ms (Figure 5C). The mean AUC-GMFA in the 2.00-8.00 ms time window was 70.8 µV*ms in the TS condition and 54.3 µV*ms in the CS+TS condition. This suggests that iTEPs evoked from stimulation of M1-LEG is sensitive to paired-pulse inhibition.

**Figure 5.**
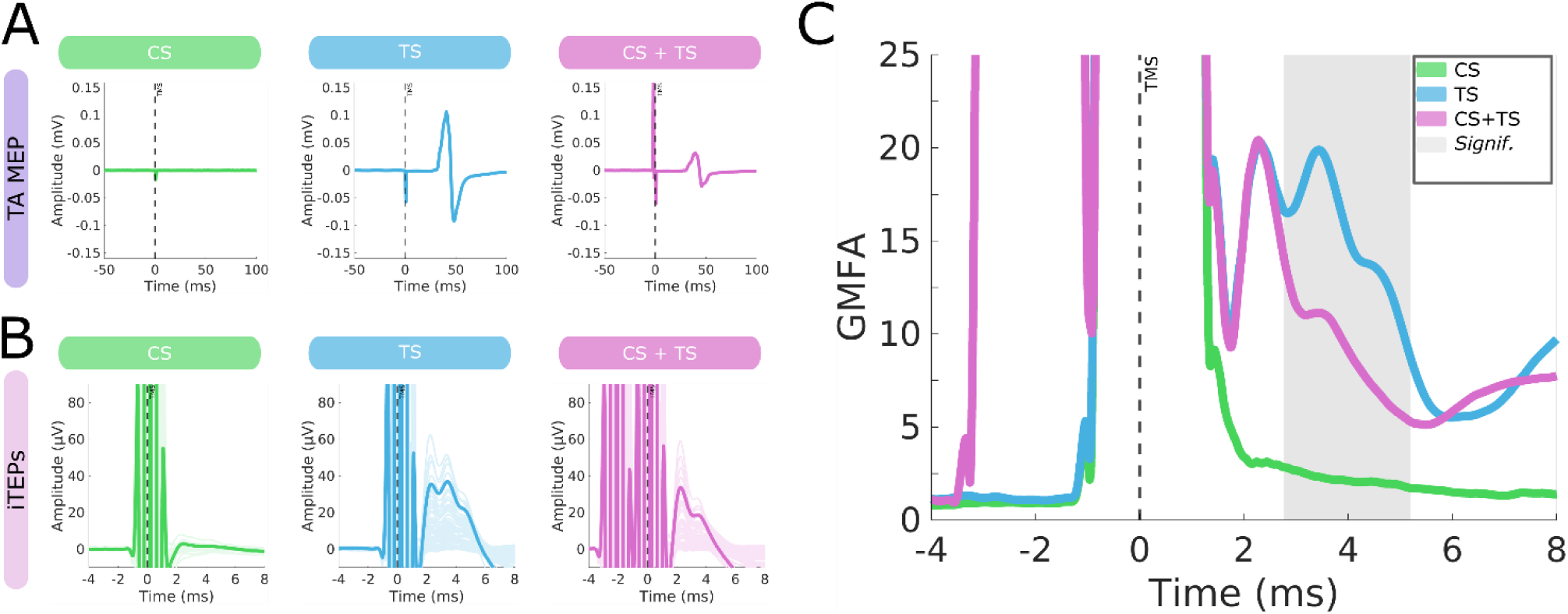
Short-latency inhibition of immediate TEPs of M1-LEG. (A) TA MEPs for CS (90% AMT, left), TS (110% RMT, middle) and CS+TS (right). (B) Butterfly plots of immediate EEG responses following stimulation of M1-LEG for CS stimulation (90% AMT, left), TS stimulation (110% RMT, middle), and CS+TS stimulation (right). The electrode highlighted in the butterfly plot represents an electrode close to the site of stimulation. iTEP responses appeared inhibited for the CS+TS condition compared to the TS condition. (C) Group mean GMFA for the CS, TS, and CS+TS condition. The shaded regions reflect significant time-window where the GMFA differed between the TS and the CS+TS condition. The CS condition is shown for visualization purposes only and was not included in the statistical comparison. Group data (*n*=10) shown as mean.

## Discussion

We found that TMS of M1-LEG elicits both iTEPs and subsequent canonical M1 TEP responses. Importantly, M1-LEG iTEPs exhibited similar physiological properties previously described for M1-HAND, including intensity dependence and sensitivity to a paired-pulse inhibition paradigm. These results provide converging evidence that iTEPs reflect motor cortical excitability and demonstrate that the cortical circuits generating immediate and subsequent TMS-evoked responses are similarly organized across upper- and lower-limb motor representations. By extending TMS-EEG characterization to M1-LEG, the present work establishes a physiological framework for investigating the cortical and corticofugal processes underlying lower-extremity motor function in both health and disease.

### Single-pulse TMS evokes iTEPs in human M1-LEG

A central finding of this study is that stimulation of M1-LEG reliably evoked immediate TMS-evoked potentials (iTEPs) with temporal and spatial characteristics that closely resemble those previously described following stimulation of M1-HAND (Beck et al., 2024a; Stango et al., 2025). Akin to M1-HAND, iTEPs evoked following M1-LEG stimulation consisted of a series of high-frequency peaks superimposed on a slower response and displayed a topographic distribution centered around the stimulated pericentral site. Following M1-LEG stimulation, response amplitudes were maximal over midline sensorimotor electrodes, consistent with a generator located in the medial sensorimotor cortex containing the cortical representation of the lower extremity muscles. This localization is well aligned with the established medial location of motor cortical areas controlling the ankle muscles (Lotze et al., 2003; Penfield and Boldrey, 1937; Terao et al., 1994). Although the motor cortical representation of the right TA muscle was targeted, we cannot exclude based on the topographical distribution of the EEG response, that bilateral activation of the medial motor cortices was achieved. Nevertheless, together with previous observations from M1-HAND, these findings suggest that iTEPs represent a reproducible physiological signature of M1 stimulation across somatotopic representations.

The similarity between M1-LEG and M1-HAND iTEPs further supports the notion that similar cortical mechanisms underlie the immediate response to TMS across somatotopic subdivisions of the motor cortex. Notably, the high-frequency components of the iTEP resemble the temporal structure of corticospinal I-wave activity, raising the possibility that they reflect physiological properties associated with densely myelinated motor cortical circuits specialized for rapid signal transmission. Consistent with this interpretation, previous studies have demonstrated that comparable iTEPs are not observed following stimulation of parietal control sites (Beck et al., 2024a), and that response amplitudes progressively decrease with increasing distance from M1 (Nuyts et al., 2025). Collectively, these observations suggest that iTEPs preferentially arise from motor cortical regions, although future studies are needed to determine whether similar immediate responses can be evoked from non-motor areas exhibiting comparable microstructural or electrophysiological characteristics.

It has been debated to which extent TMS-evoked scalp muscle activity could contribute to iTEP (Stango et al., 2025). The present findings also argue against scalp muscle activity as an important contributing source of the iTEP. Stimulation of M1-LEG was performed close to the vertex, a region largely devoid of cranial musculature and commonly regarded as a favorable target for avoiding TMS-related muscle artifacts (Mutanen et al., 2013). The robust presence of iTEPs under these conditions therefore strengthens the interpretation that these responses primarily reflect cortical activation rather than peripheral contamination. From a methodological perspective, the medial motor cortex may consequently represent a particularly attractive target for studying human motor cortical physiology with TMS-EEG, offering reliable access to immediate and canonical cortical responses while minimizing the likelihood of confounding scalp muscle artifacts.

### Intensity dependence of iTEPs elicited by TMS of M1-LEG

A second key finding is that M1-LEG iTEPs exhibit a clear stimulation intensity–response relationship. Immediate TEPs emerged around active motor threshold and were hence detectable at stimulation intensities below those required to elicit MEPs at rest. In addition, response amplitudes systematically increased with stimulation intensity. This recruitment profile closely resembles that reported for iTEPs elicited from M1-HAND (Beck et al., 2024a; Christiansen, Song et al., 2026) and shares important features with the recruitment characteristics of corticospinal volleys recorded epidurally following stimulation of both upper- and lower-extremity motor representations (Di Lazzaro et al., 1998b; V Di Lazzaro et al., 2001; V. Di Lazzaro et al., 2001). These observations further support the notion that iTEPs provide a sensitive electrophysiological marker of cortical recruitment by TMS.

Stimulation of M1-LEG requires higher stimulation intensities than stimulation of M1-HAND, largely because the lower-extremity representation lies deep within the interhemispheric fissure (Lotze et al., 2003). The MEP threshold for TA is thought to be lower than in other leg muscles in part due to the larger expression of monosynaptic links to the spinal motoneurons (Brouwer and Ashby, 1992; Petersen et al., 2003). This favors TA over other leg muscles as the target for TMS-EEG studies of M1-LEG. Given the depth-focality trade-off of TMS (Deng et al., 2013) effective stimulation of M1-LEG is achieved at the expense of reduced focality, increasing the likelihood of co-activating adjacent motor cortical regions. The iTEPs observed in the present study should therefore not be interpreted as arising exclusively from the tibialis anterior representation. Instead, they likely reflect the summed activity of neuronal populations within the medial motor cortex, potentially including representations of other lower-extremity, axial, and proximal upper extremity musculature (Ferbert et al., 1992). This broader recruitment may contribute to the larger iTEP amplitudes observed following M1-LEG stimulation relative to those previously reported for M1-HAND despite broadly similar peak latencies and temporal structure (Beck et al., 2024a; Christiansen, Song et al., 2026; Nuyts et al., 2026, 2025). These findings also highlight an important distinction between iTEPs and downstream MEPs. Whereas MEPs primarily reflect corticospinal output to a specific muscle, iTEPs appear to index cortical recruitment at the level of local and distributed neuronal populations. Accordingly, caution is warranted when directly linking individual iTEP components to the activation of a particular effector or motoneuronal pool. Although stimulation was targeted to the optimized tibialis anterior hotspot, the extent to which the recorded iTEPs reflect activity specifically associated with corticospinal projections to the tibialis anterior remains uncertain and will require further investigation using approaches with greater spatial specificity. In addition, cortical and corticospinal responses may diverge evident as differential impact on iTEPs and MEPs in cases where state-dependent changes in spinal and cortical circuits oppose (Nuyts et al., 2026). Accordingly, iTEPs and MEPs should be viewed as complementary measures indexing distinct levels of motor system excitability.

### Short-latency intracortical inhibition of iTEPs evoked by TMS of M1-LEG

A third main finding is that both iTEPs and MEPs elicited by M1-LEG stimulation were consistently suppressed by a subthreshold conditioning pulse given 2.3 ms before the test pulse, demonstrating robust SICI. The effects of SICI on iTEPs were restricted to the second and third peak of the iTEP, while the first iTEP peak was not modulated by the preceding conditioning response. Although the magnitude of short-latency inhibition varied between participants, suppression of the second iTEP peak was consistently observed across all individuals, indicating that the immediate cortical response is sensitive to intracortical inhibitory mechanisms. This finding extends recent observations from M1-HAND, where paired-pulse stimulation similarly attenuated the amplitude of the second iTEP peak (Christiansen, Song et al., 2026). The presence of short-latency paired-pulse inhibition in both upper-and lower-extremity motor representations supports the notion that iTEPs reflect fundamental physiological processes involved in motor cortex excitability.

The observed suppression is also consistent with evidence from epidural recordings demonstrating attenuation of indirect corticospinal volleys following paired-pulse stimulation of both M1-HAND and M1-LEG (Di Lazzaro et al., 1998b; V. Di Lazzaro et al., 2001). These epidural electrophysiological studies attributed the reduction in corticospinal output primarily to the suppression of later I-waves, suggesting that intracortical inhibitory circuits modulate corticospinal neuron activity before descending signals enter the corticospinal tract. The close correspondence between paired-pulse effects on iTEPs, MEPs, and epidurally recorded corticospinal volleys therefore provides additional support for the hypothesis that iTEPs are closely linked to the physiological processes underlying corticospinal recruitment.

Nevertheless, the relationship between individual iTEP components and specific corticospinal volleys remains unresolved. While paired-pulse inhibition of M1-HAND has been linked to suppression of multiple late I-waves, invasive recordings obtained following M1-LEG stimulation have primarily demonstrated inhibition of the third and fourth I-waves (Di Lazzaro et al., 2001). Whether specific iTEP peaks correspond to particular components of the corticospinal volley is therefore currently unknown. The partial convergence, but not complete correspondence, between epidural and EEG findings raises the possibility that iTEPs capture aspects of local cortical inhibition that are not directly reflected in downstream corticospinal recordings.

The present analyses also do not allow firm conclusions regarding which features of the iTEP complex are preferentially suppressed by intracortical inhibition. Visual inspection suggests attenuation of both the high-frequency peaks and the slower underlying component, but it remains challenging to separate the relative contribution of these signal components and still resolve single iTEP peaks (Christiansen, Song et al., 2026). Future studies should therefore seek to disentangle the fast and slow elements of the iTEP response while preserving sensitivity to individual peaks. Such work may provide further insights into the cortical circuits responsible for generating immediate TMS-evoked activity and their modulation by inhibitory interneuron networks.

### Early and late TEPs evoked by TMS of the M1-LEG

In addition to the iTEPs, stimulation of M1-LEG elicited a sequence of early and late TEP responses that closely resembled the canonical motor-cortical response previously described following M1-HAND stimulation (Beck et al., 2024b). The temporal organization of these responses, including the N15, P30, N45, P60, and N100, was largely preserved over both hand and leg motor representations, supporting the notion that common network mechanisms contribute to cortical processing of TMS within M1. However, an important difference emerged in the spatial distribution of the early responses. Following M1-LEG stimulation, the N15, P30, N45, P60, and N100 responses all exhibited topographies consistent with generators located in the medial sensorimotor cortex, whereas following M1-HAND stimulation only the N100 exhibits a predominantly central topography, likely reflecting a residual sensory confound (Conde et al., 2019). This observation is consistent with the distinct anatomical location of the leg representation within the interhemispheric fissure and further supports the site-specific origin of the early TEP components.

The present findings therefore also align with the prevailing view that early TEP components are thought to predominantly reflect reverberant local cortical activation and regional network dynamics, whereas later responses are increasingly influenced by widespread cortical processing as well as sensory co-stimulation arising from the TMS pulse (Hernandez-Pavon et al., 2023; Massimini et al., 2026). The preservation of a medial topography for several early components following M1-LEG stimulation therefore provides further evidence that these responses retain information about the stimulated cortical region. In contrast, the broader distribution of later components likely reflects the contribution of shared large-scale cortical networks and sensory-evoked activity that is relatively independent of stimulation site.

To our knowledge, this is the first study to comprehensively characterize TEPs from the leg motor cortex at latencies below 100 ms.. Previous TMS-EEG studies targeting M1-LEG have largely focused on later evoked responses or post-stimulation oscillatory dynamics (Fecchio et al., 2017; Huntington et al., 2026; Massimini et al., 2007). By extending the characterization of TEPs into the immediate and early post-stimulus period, the present results demonstrate that the leg motor cortex exhibits a motor-typical sequence of local and distributed network responses that closely parallels those observed in the hand representation, while retaining topographic features consistent with its distinct anatomical location within the motor homunculus.

## Conclusion and perspectives

The present study demonstrates for the first time that TMS of the leg representation of primary motor cortex elicits robust immediate cortical potentials, preceding the canonical sequence of later TMS-evoked responses. M1-LEG iTEPs share the defining physiological features of hand-area iTEPs, including threshold dependence, intensity scaling, and sensitivity to short-interval intracortical inhibition. These findings establish that iTEPs are not unique to M1-HAND but represent a general physiological response of human motor cortex.

The striking similarity between M1-LEG and M1-HAND iTEPs suggests that the mechanisms generating immediate cortical responses are conserved across somatotopically distinct motor representations. Moreover, the intensity dependence and paired-pulse sensitivity of M1-LEG iTEPs provide further support for their proposed relationship to the cortical generators of corticospinal I-waves and associated inhibitory circuits.

Beyond their physiological significance, M1-LEG iTEPs may have important clinical utility (Ziemann et al., 2026). Disorders affecting lower-extremity motor function, including stroke, spinal cord injury, multiple sclerosis, and other neurological diseases, are frequently associated with elevated motor thresholds, impaired corticospinal conduction, or absent motor-evoked potentials. In these situations, conventional TMS measures provide limited information about residual cortical function. Because iTEPs directly index cortical responses to TMS, they may offer a sensitive, non-invasive measure of motor cortical excitability and inhibition even when peripheral motor responses cannot be recorded reliably.

By extending iTEPs to the leg motor cortex, the present work broadens the applicability of TMS-EEG for probing excitatory and inhibitory circuit dynamics throughout the human motor system. Future studies should establish the reliability, sensitivity, and clinical relevance of M1-LEG iTEPs and determine their potential as biomarkers of cortical function, plasticity, and recovery in neurological disorders affecting lower-limb motor control.

## Acknowledgements

The authors would like to thank Ditte Haagerup and Agata Banach for their help with data acquisition.

## Author contributions

MMB, MN, JR, HR & LC conceptualized and designed the study. MMB, MN & LS collected data. MMB analyzed data. MMB & LC drafted the first version of manuscript. All authors have critically reviewed the manuscript and approved of the final version.

## Supplementary information

In table S1, peak latencies and amplitudes of the GMFA are extracted per subject

**Table S1:** Individual latencies and amplitudes of the GMFA for the 110% RMT condition.

| *Subject* | **Peak 1** |  | **Peak 2** |  | **Peak 3** |  |
| --- | --- | --- | --- | --- | --- | --- |
|  | *Latency (ms)* | *Amplitude* | *Latency (ms)* | *Amplitude* | *Latency (ms)* | *Amplitude* |
| #1 | 2,26 | 16,40 | 3,56 | 20,06 | 4,92 | 18,57 |
| #2 | 2,44 | 25,25 | — | — | — | — |
| #3 | 2,18 | 13,38 | — | — | 4,42 | 2,47 |
| #4 | 2,26 | 19,99 | 3,52 | 24,73 | 4,60 | 24,51 |
| #5 | 2,34 | 5,78 | 3,78 | 8,48 | 5,20 | 10,22 |
| #6 | — | — | 3,54 | 31,64 | — | — |
| #7 | 2,28 | 25,80 | 3,32 | 24,92 | 4,44 | 18,76 |
| #8 | 2,18 | 25,71 | 3,24 | 24,23 | — | — |
| #9 | 2,52 | 16,63 | — | — | — | — |
| #10 | — | — | 3,40 | 20,18 | 4,50 | 18,88 |
| #11 | 2,28 | 26,88 | 3,46 | 22,65 | — | — |
| #12 | 2,06 | 35,06 | 3,34 | 22,25 | — | — |
| #13 | 2,42 | 19,36 | — | — | — | — |
| #14 | 2,28 | 31,87 | 3,34 | 26,65 | — | — |
| #15 | 2,26 | 13,12 | 3,54 | 15,45 | 4,84 | 12,57 |
| #16 | 2,32 | 25,56 | 3,56 | 28,68 | — | — |
| Mean | 2,29 | 21,49 | 3,47 | 22,50 | 4,70 | 15,14 |
| Median | 2,28 | 22,63 | 3,49 | 23,44 | 4,60 | 18,57 |
| SD | 0,12 | 7,98 | 0,15 | 6,11 | 0,29 | 7,28 |

